# Pirfenidone improves adipose dysfunction and obesity-driven steatohepatitis via mTORC1 signaling

**DOI:** 10.64898/2026.03.20.713092

**Authors:** Yu Seol Lee, Ji Yun Bang, Da Hyun Lee, Da Ye Kim, Shin Young Cha, Eo Jin Lee, Jisu Han, Soo Han Bae

**Affiliations:** Department of Biomedical Sciences, Yonsei University College of Medicine, Seoul, Republic of Korea; Graduate School of Medical Science, BK21 PLUS Project, Yonsei University College of Medicine, Seoul 03722, Republic of Korea

**Keywords:** Pirfenidone, mTORC1 signaling, Adipose tissue dysfunction, Adipose tissue remodel, VAT fibrosis, Adipose tissue–liver crosstalk, MASLD, MASH

## Abstract

Obesity-driven metabolic dysfunction–associated steatotic liver disease (MASLD) and metabolic dysfunction–associated steatohepatitis (MASH) are shaped by depot-specific adipose tissue dysfunction, including maladaptive expansion and visceral adipose tissue (VAT) fibrosis. Pirfenidone, an anti-fibrotic agent, improves experimental liver disease. However, its actions on adipose depots and adipose–liver crosstalk remain unclear. Here, we identify pirfenidone as a modulator of mechanistic target of rapamycin complex 1 (mTORC1)-dependent adipose tissue remodeling with divergent outputs in subcutaneous and visceral fat. In diet-induced obese MASH mice, pirfenidone decreased subcutaneous adipose tissue (SAT), inhibiting mTORC1-driven lipogenesis and enhancing oxidative lipid metabolism. Pirfenidone attenuated VAT fibrosis by suppressing an mTORC1–mothers against decapentaplegic homolog 3 (SMAD3)–yes-associated protein (YAP) axis and extracellular matrix gene programs. Pirfenidone also lowered hepatic triglycerides, improved steatosis and fibrosis, reduced hepatic mTORC1 activity. Conditioned medium from fibrotic adipocytes induced lipogenic, inflammatory, and pro-fibrotic programs in AML12, which effects that were blunted by pirfenidone. These data reveal adipose tissue-centered actions of pirfenidone that link mTORC1 remodeling to improved obesity-associated liver disease.

## Introduction

Metabolic dysfunction–associated steatotic liver disease (MASLD) and its progressive inflammatory form, metabolic dysfunction–associated steatohepatitis (MASH), are rapidly increasing in parallel with the global obesity epidemic (Eslam *et al*, 2020; Younossi *et al*, 2023). Growing evidence suggests that MASLD/MASH result from systemic metabolic dysfunction and inter-organ crosstalk rather than a liver-intrinsic process (Steinberg *et al*, 2025), and adipose tissue functions as an active metabolic and endocrine organ that shapes systemic lipid flux and inflammatory signaling (Mitrou *et al*, 2010). Therefore, defining the signaling pathways that govern adipose-to-liver crosstalk is essential for understanding disease progression and identifying therapeutic entry points (Friedman *et al*, 2018; Tilg & Moschen, 2010).

In obesity, adipose tissue expands to buffer excess nutrients. However, this adaptive response can become pathological, leading to adipocyte hypertrophy, impaired lipid storage, altered cytokine/chemokine secretion, inflammatory activation, and ectopic lipid delivery to the liver (Azzu *et al*, 2020; Frayn, 2002). Adipose tissue dysfunction is depot-specific: subcutaneous adipose tissue (SAT) and visceral adipose tissue (VAT) exhibit distinct remodeling programs and metabolic consequences (Ibrahim, 2010; Lee *et al*, 2013). VAT expansion is frequently accompanied by excessive extracellular matrix deposition and fibrosis, which increase tissue stiffness, impair adipocyte plasticity, and are associated with metabolically unhealthy obesity (Beals *et al*, 2021; Marcelin *et al*, 2022). The signaling logic that drives depot-selective adipose tissue remodeling and transmits adipose tissue-derived cues to hepatic lipotoxicity, inflammation, and fibrogenesis is not yet defined (Bilson *et al*, 2024; Sun *et al*, 2014).

Mechanistic target of rapamycin complex 1 (mTORC1) is a nutrient-sensing hub that integrates growth factors, amino acids, and energy status to control anabolic metabolism and cellular remodeling (Saxton & Sabatini, 2017). In adipocytes, mTORC1 promotes lipogenesis and influences adipose expansion, stress responses, and secretory programs, positioning it as a potential coordinator of obesity-induced adipose tissue remodeling (Chimin *et al*, 2017; Liu *et al*, 2016). However, the downstream outputs of mTORC1 signaling may differ among adipose depots, potentially reflecting differences in stromal composition, mechanical and pro-fibrotic microenvironments that shape extracellular matrix (ECM) accumulation and tissue architecture (Divoux *et al*, 2010; Grandl *et al*, 2016). Because adipose remodeling strongly influences lipid flux and inflammatory signaling to the liver, depot-selective mTORC1 outputs may have distinct consequences for adipose–liver crosstalk and metabolic liver disease. While mTORC1 has been linked to hepatic lipid metabolism and inflammatory signaling, (Gosis *et al*, 2022; Marcondes-de-Castro *et al*, 2023) it remains unclear whether clinically relevant anti-fibrotic agents modulate mTORC1 signaling across metabolic tissues and whether such target engagement improves adipose tissue–liver crosstalk.

Pirfenidone, which is an orally available anti-fibrotic drug approved for idiopathic pulmonary fibrosis, has anti-inflammatory and anti-fibrotic activities in experimental settings (Aimo *et al*, 2020; Behr *et al*, 2017; Torre *et al*, 2024). In models of metabolic liver disease, pirfenidone has been reported to reduce hepatic inflammation and fibrosis, suggesting a possible use beyond pulmonary fibrosis (Chen *et al*, 2019; Xi *et al*, 2021). However, despite reported hepatic benefits, whether pirfenidone alters mTORC1 activity in metabolic tissues and whether these changes reprogram depot-specific adipose remodeling—and in turn reshape adipose tissue-to-liver communication—has not been determined.

Here, we investigated whether pirfenidone modulates mTORC1-dependent remodeling in metabolic tissues and how this relates to obesity-associated liver disease. Using diet-induced obese mice with MASLD/MASH features, we show that pirfenidone suppresses mTORC1 activity in SAT, VAT, and liver, but the remodeling outputs differ among adipose tissue depots. Pirfenidone limits SAT expansion by reducing mTORC1-driven lipogenic programs while enhancing oxidative lipid metabolism-associated genes. In contrast, pirfenidone attenuates VAT fibrosis by suppressing the mTORC1–mothers against decapentaplegic homolog 3 (SMAD3)–yes-associated protein (YAP) axis and ECM gene programs, restoring adipose tissue architecture. These depot-specific improvements coincide with reduced hepatic triglyceride accumulation and amelioration of steatosis, inflammation, and fibrosis. To functionally link adipose remodeling to hepatocyte responses, we established an adipose tissue-to-liver signaling axis using a fibrotic adipocyte-conditioned medium system. Conditioned medium–based co-culture in alpha mouse liver 12 (AML12) hepatocytes induced lipogenic, inflammatory, and pro-fibrotic programs, which were blunted by pirfenidone. Together, our findings demonstrate the adipose-centered actions of pirfenidone and identify depot-selective mTORC1 remodeling as a mechanistic bridge between adipose dysfunction and obesity-associated liver disease.

## Results

### Pirfenidone improves systemic metabolic parameters and remodels adipose tissue distribution in obesity-associated MASLD

To evaluate the metabolic effects of pirfenidone in obesity-associated MASLD, we treated diet-induced obese mice with pirfenidone and assessed metabolic parameters (Fig. 1A). Pirfenidone had minimal effects on body weight in normal diet-fed mice, whereas it significantly reduced body weight in high-fat diet (HFD)-fed mice, indicating a diet-dependent impact of pirfenidone under obese conditions (Fig. 1B). Consistent with these results, gross morphology and tissue weights showed depot-selective changes in adiposity. Pirfenidone reduced inguinal white adipose tissue (iWAT) and liver weights, whereas epididymal white adipose tissue (eWAT) and brown adipose tissue (BAT) weights were largely unchanged (Fig. 1C,D). Systemically, pirfenidone reduced the levels of circulating non-esterified fatty acids (Fig. 1E) and of serum alanine aminotransferase (Fig. 1F). Functionally, pirfenidone improved glucose homeostasis, as shown by improved glucose tolerance during oral glucose tolerance tests (Fig. 1G,H) and enhanced insulin responsiveness during insulin tolerance tests (Fig. 1I,J). Together, these data indicate that pirfenidone reduces systemic metabolic stress and markers of liver injury and induces depot-selective adipose remodeling in obese MASLD mice.

**Figure 1.**
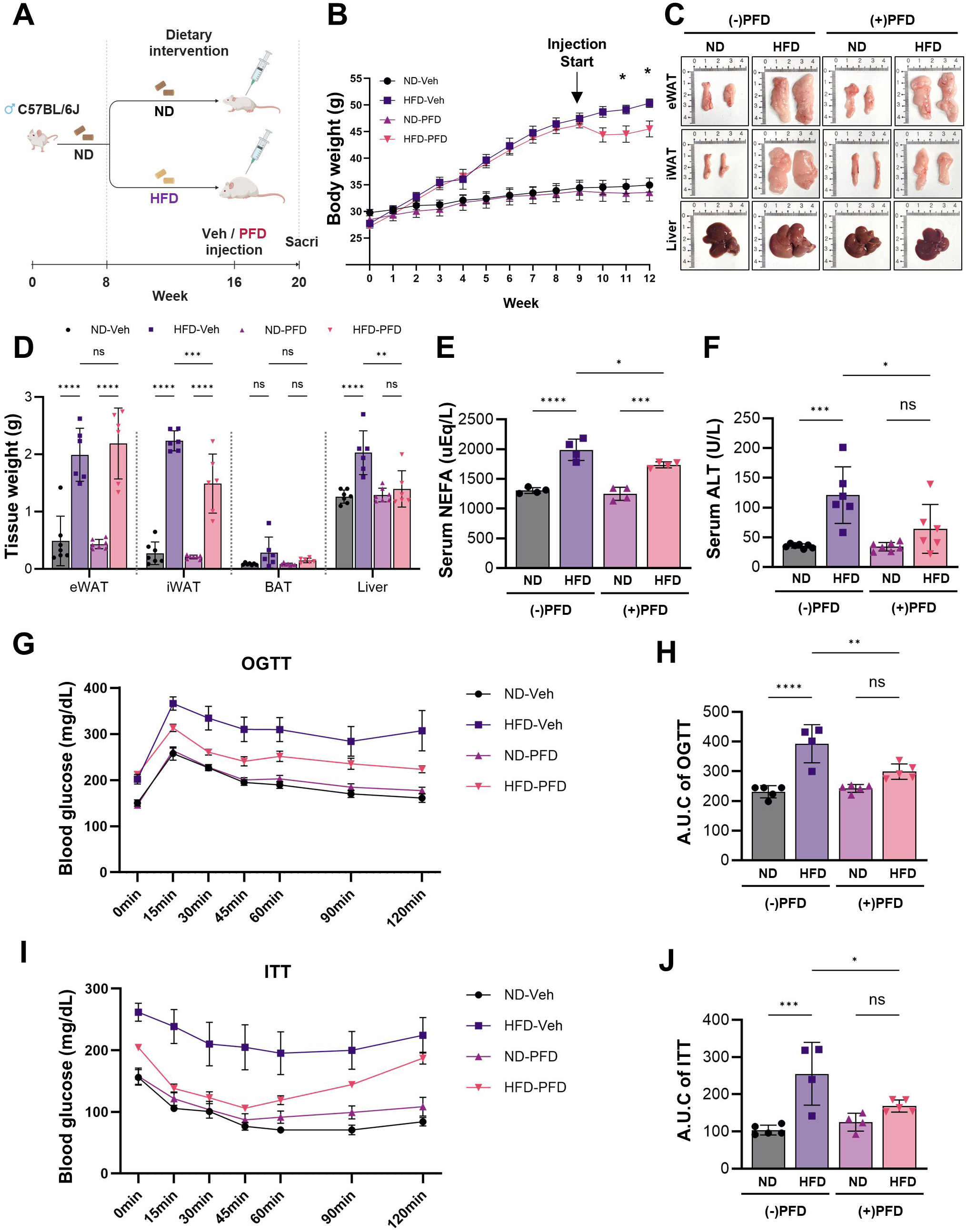
Pirfenidone improves the systemic metabolic profile and remodels adipose tissue distribution in obese MASLD mice. **(A)** Experimental scheme of the diet-induced obese metabolic dysfunction–associated steatotic liver disease (MASLD) model. Male C57BL/6J mice were fed a high-fat diet (HFD) and treated with pirfenidone (PFD) or vehicle after 9-weeks of HFD-feeding. **(B)** Body weight trajectories of mice fed normal diet (ND) or HFD with vehicle or PFD treatment. **(C)** Representative gross morphologies of adipose tissue depots (eWAT, iWAT, and BAT) and liver from ND- and HFD-fed mice treated with vehicle or PFD. **(D)** Tissue weights for epididymal white adipose tissue (eWAT), inguinal white adipose tissue (iWAT), brown adipose tissue (BAT), and liver. **(E)** Serum non-esterified fatty acid (NEFA) levels. **(F)** Serum alanine aminotransferase (ALT) levels. **(G)** Oral glucose tolerance test (OGTT) curves. **(H)** Area under the curve (AUC) analysis of the OGTT results. **(I)** Insulin tolerance test (ITT) curves. **(J)** AUC analysis of the ITT results. Data are presented as the mean ± SEM. Each dot represents an individual mouse. Statistical significance was determined by one-way or two-way ANOVA with Tukey’s multiple-comparison test, as appropriate. *P < 0.05, **P < 0.01, ***P < 0.001; ns, not significant.

### Pirfenidone promotes oxidative remodeling and browning programs in inguinal white adipose tissue

Given the depot-selective reduction of iWAT mass in pirfenidone-treated obese mice (Fig. 1), we next examined whether pirfenidone reshapes adipocyte morphology and metabolic programs in iWAT. Histological analysis showed that HFD feeding induced marked adipocyte hypertrophy, which was attenuated by pirfenidone (Fig. 2A). Quantification of adipocyte size distribution confirmed a leftward shift in adipocyte diameter with pirfenidone treatment, indicating reduced hypertrophic expansion (Fig. 2B). Consistent with decreased lipid storage, pirfenidone reduced protein levels of adipose differentiation-related protein (ADRP)/perilipin 2 (PLIN2) in iWAT (Fig. 2C). At the transcriptional level, pirfenidone suppressed the lipogenic gene program induced by HFD (Fig. 2D), supporting a reduction in de novo lipid synthesis. In parallel, pirfenidone further increased expression of fatty acid oxidation–associated genes in HFD-fed mice (Fig. 2E), consistent with enhanced lipid utilization. Notably, pirfenidone also upregulated browning/thermogenic gene signatures and increased the level of peroxisome proliferator-activated receptor-γ coactivator 1α (PGC1α) in iWAT (Fig. 2F,G). Finally, inflammatory gene expression in iWAT was reduced by pirfenidone (Fig. 2H), consistent with an overall improvement of adipose tissue remodeling. Together, these data indicate that under obese conditions, pirfenidone reprograms iWAT toward an oxidative, lipid-utilizing state, which is characterized by reduced adipocyte hypertrophy and lipid storage, induction of browning-associated signatures, and decreased expression of inflammatory genes.

**Figure 2.**
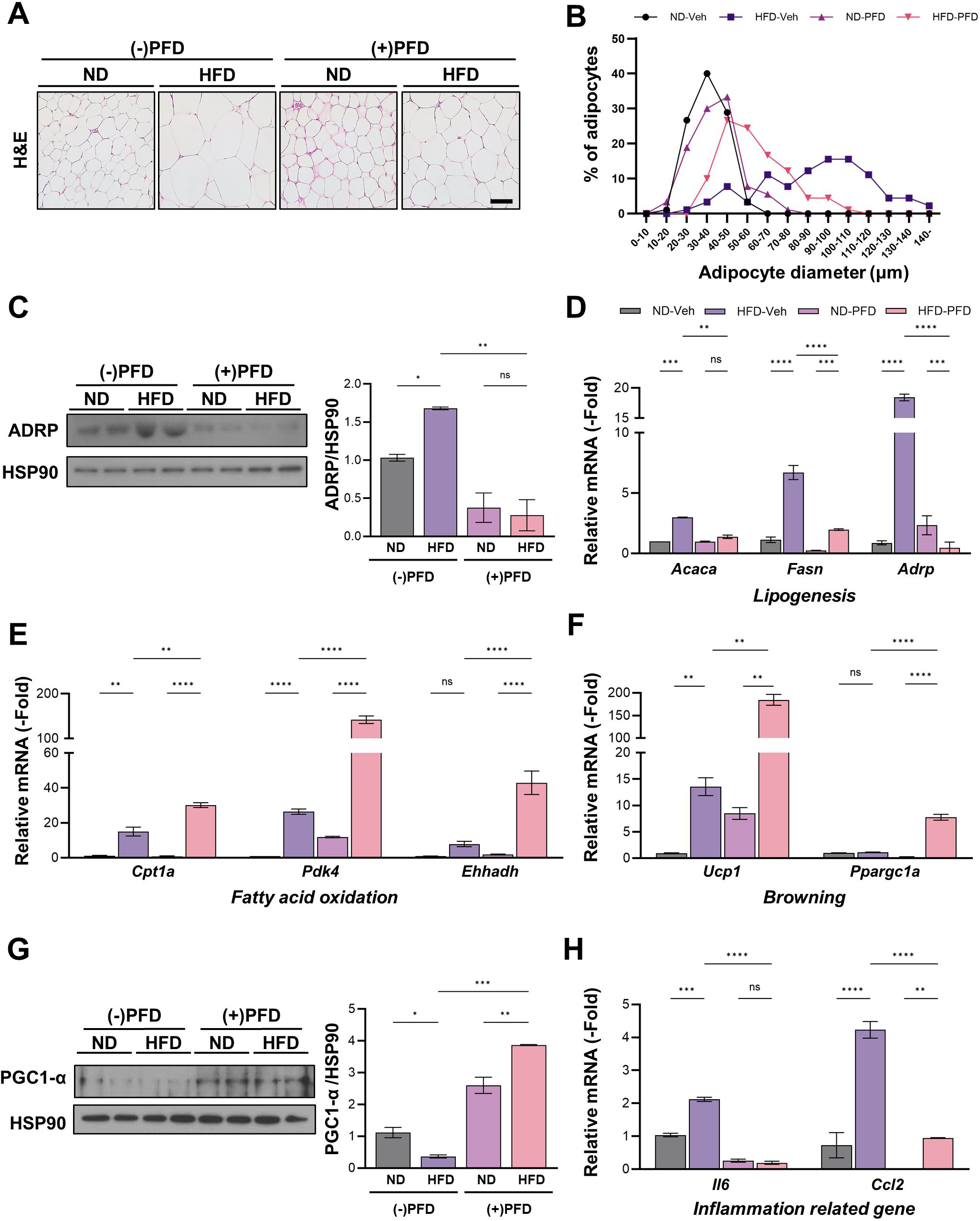
Pirfenidone modulates lipid metabolism in inguinal white adipose tissue. **(A)** Representative H&E staining of inguinal white adipose tissue (iWAT) from normal diet (ND)- and high-fat diet (HFD)-fed mice treated with vehicle or pirfenidone. Scale bar, 100 μm. **(B)** Distribution of adipocyte diameters in iWAT from ND- and HFD-fed mice with or without pirfenidone treatment. **(C)** Immunoblot analysis of adipose differentiation-related protein (ADRP) in iWAT, with quantification normalized to HSP90. **(D)** Relative mRNA expression levels of lipogenesis-related genes (*Acaca, Fasn,* and *Adrp*) in iWAT. **(E)** Relative mRNA expression levels of fatty acid oxidation–related genes (*Cpt1a, Pdk4,* and *Ehhadh*) in iWAT. **(F)** Relative mRNA expression levels of browning-associated genes (*Ucp1* and *Ppargc1a*) in iWAT. **(G)** Immunoblot analysis of PGC1α protein levels in iWAT, with quantification normalized to HSP90. **(H)** Relative mRNA expression levels of inflammation-related genes (*Il6* and *Ccl2*) in iWAT. Data are presented as the mean ± SEM (n = 5 mice per group). Statistical significance was determined by two-way ANOVA followed by Tukey’s multiple-comparison test. *P < 0.05; **P < 0.01; ***P < 0.001; ****P < 0.0001; ns, not significant.

### Pirfenidone suppresses adipocyte lipid accumulation by inhibiting mTORC1-dependent lipogenic and lipid-uptake programs in vitro

To mechanistically examine the adipose remodeling observed in vivo, we assessed the effects of pirfenidone during 3T3-L1 adipocyte differentiation using stage-specific treatment phases (Fig. 3A). Oil Red O staining showed that pirfenidone reduced neutral lipid accumulation in differentiated adipocytes, with the magnitude depending on the timing of exposure (Fig. 3B,C). Consistent with this phenotype, pirfenidone decreased expression of genes involved in de novo lipogenesis, lipid droplet storage, and fatty acid uptake (Fig. 3D). In contrast, transcripts associated with fatty acid oxidation were largely unchanged, suggesting that pirfenidone limits lipid loading rather than directly enhancing oxidative disposal under these conditions. Immunoblotting confirmed suppression of the adipogenic program, as reflected by reduced peroxisome proliferator-activated receptor-γ (PPARγ) levels (Fig. 3E,F). Notably, pirfenidone decreased mTORC1 activity, as evidenced by reduced phosphorylation of S6 kinase 1 (S6K1) and S6 (Fig. 3E,G,H). Consistently, pirfenidone-treated adipocytes exhibited reduced adenosine monophosphate-activated protein kinase (AMPK) phosphorylation (Fig. 3E), suggesting that pirfenidone primarily modulates mTORC1 signaling under these differentiation conditions. Taken together, these data support a model in which pirfenidone restrains adipocyte lipid accumulation by dampening mTORC1-dependent lipogenic and lipid-uptake programs.

**Figure 3.**
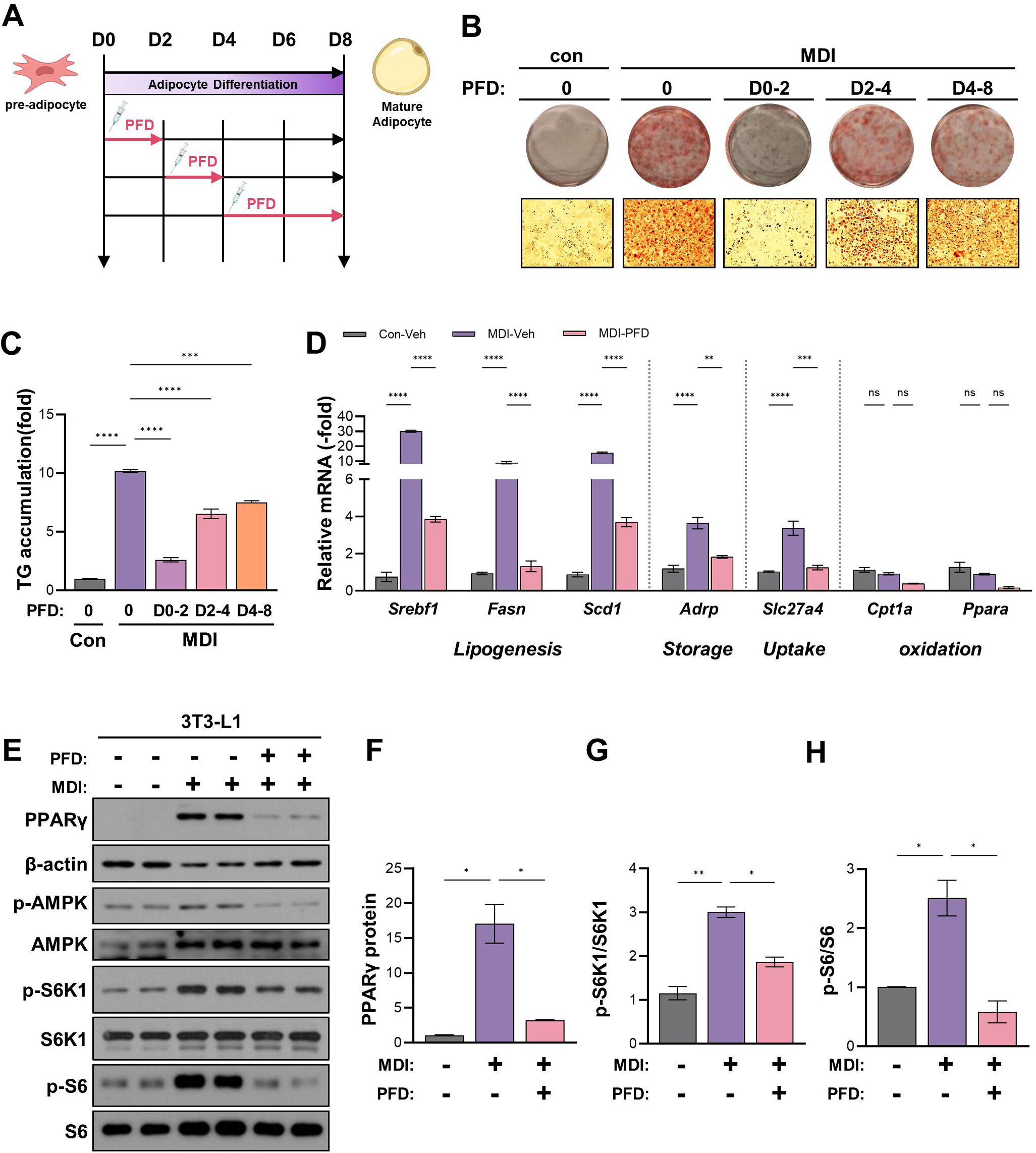
Pirfenidone attenuates adipocyte hypertrophy by inhibiting lipogenesis and lipid uptake via mTORC1 signaling in vitro. **(A)** Schematic of the 3T3-L1 adipocyte maturation protocol and timing of pirfenidone (PFD) treatment during adipogenesis. **(B)** Representative Oil Red O staining of differentiated adipocytes treated with PFD during the indicated differentiation windows (D0–2, D2–4, D4–8). **(C)** Quantification of Oil Red O staining intensity in control (Con), MDI-treated, and MDI plus PFD-treated adipocytes. **(D)** Relative mRNA expression levels of genes involved in lipogenesis (*Srebf1, Fasn,* and *Scd1*), lipid storage (*Adrp*), lipid uptake (*Slc27a4*), and fatty acid oxidation (*Cpt1a* and *Ppara*) in differentiated 3T3-L1 adipocytes. **(E)** Immunoblot analysis of PPARγ, phosphorylated AMPK (p-AMPK), total AMPK, phosphorylated p70S6 kinase (p-S6K1), total S6K1, phosphorylated S6 (p-S6), and total S6 in 3T3-L1 adipocytes treated with MDI and/or PFD. β-actin was used as a loading control. **(F)** Quantification of PPARγ protein levels normalized to β-actin. **(G)** Quantification of p-S6K1 levels normalized to total S6K1. **(H)** Quantification of p-S6 levels normalized to total S6. Data are presented as the mean ± SEM. Statistical significance was determined by one-way ANOVA with Tukey’s multiple-comparison test. *P < 0.05, **P < 0.01, ***P < 0.001; ns, not significant. MDI denotes a differentiation cocktail consisting of 3-isobutyl-1-methylxanthine (IBMX, 520 µM), dexamethasone (1 µM), and insulin (1 µg/mL).

### Pirfenidone suppresses fibro-inflammatory remodeling in epididymal adipose tissue

Although pirfenidone did not significantly change eWAT mass in obese mice (Fig. 1D), we examined whether it improves the quality of VAT remodeling. Masson’s trichrome and Sirius Red staining showed extensive ECM deposition in eWAT from HFD-fed mice, which was reduced by pirfenidone (Fig. 4A,B). Similarly, pirfenidone blunted fibrosis-associated protein signatures, including α-smooth muscle actin (α-SMA), fibronectin (FN), and collagen VI (Fig. 4C,D). At the transcriptional level, pirfenidone attenuated pro-fibrotic gene programs, including collagens (*Col1a1, Col3a1*) and additional matrix-associated genes (*Fn1, Serpine1, Ccn2, and Tgfb1*) (Fig. 4E). In parallel, the expression levels of the inflammatory mediators *Tnf* and *Ccl2* were decreased (Fig. 4F). Consistent with these molecular changes, hematoxylin and eosin (H&E) and F4/80 staining showed reduced adipocyte hypertrophy and diminished macrophage accumulation in eWAT from pirfenidone-treated mice (Fig. 4G–I). Collectively, these findings indicate that pirfenidone alleviates visceral adipose fibrosis and inflammation, improving fibro-inflammatory remodeling without a detectable change in eWAT mass.

**Figure 4.**
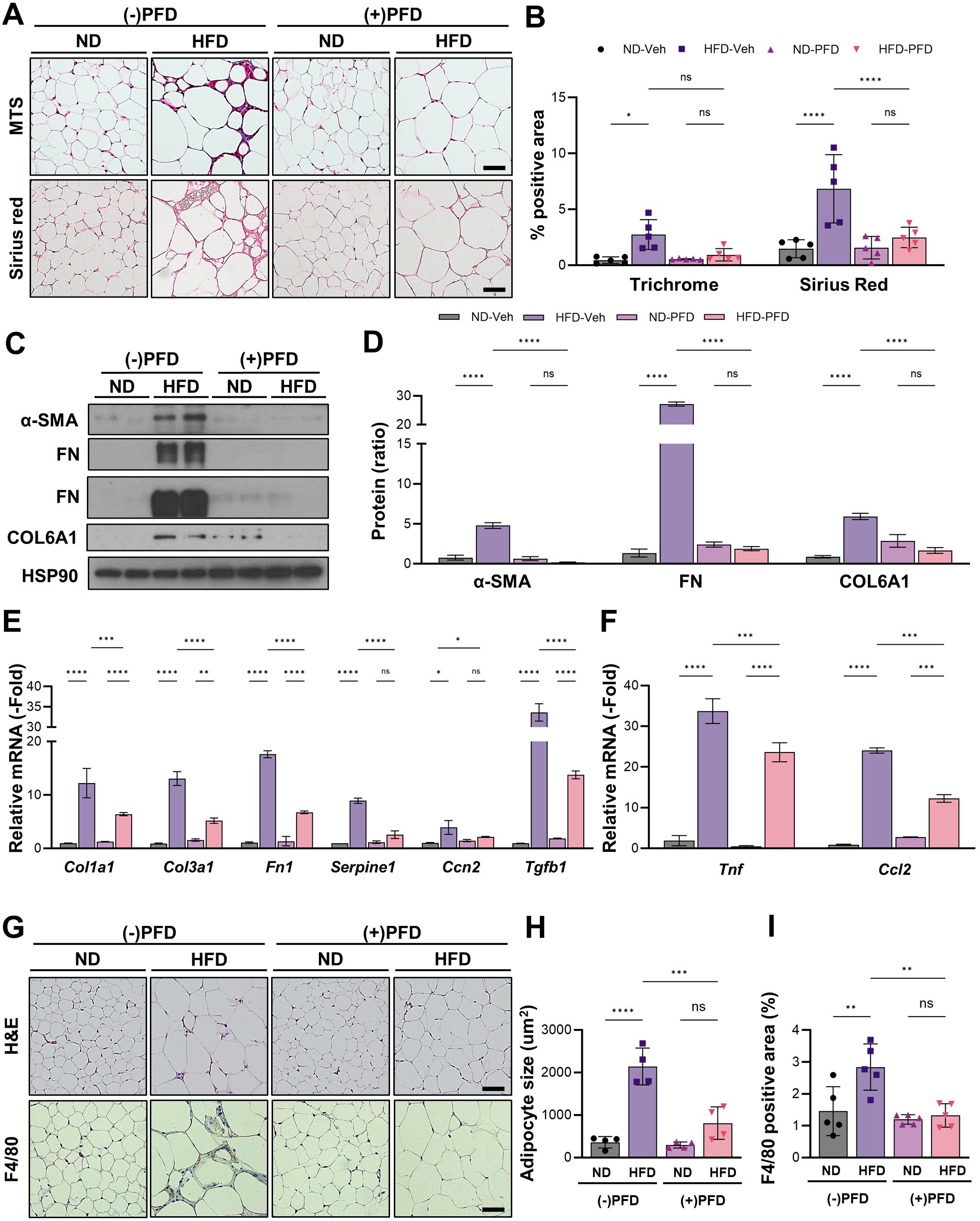
Pirfenidone suppresses fibrosis and inflammation in epididymal adipose tissue. **(A)** Representative Masson’s trichrome and Sirius Red staining of epididymal white adipose tissue (eWAT) from normal diet (ND)- and high-fat diet (HFD)-fed mice treated with vehicle or pirfenidone. Scale bars, 100 μm. **(B)** Quantification of fibrotic area based on Masson’s trichrome and Sirius Red staining in eWAT. **(C)** Immunoblot analysis of fibrosis-associated proteins α-smooth muscle actin (α-SMA), fibronectin (FN), and collagen VI α1 (Col6A1) in eWAT, with HSP90 as a loading control. **(D)** Quantification of α-SMA, FN, and COL6A1 protein levels normalized to HSP90. **(E)** Relative mRNA expression levels of extracellular matrix and fibrosis-related genes (*Col1a1, Col3a1, Fn1, Serpine1, Ccn2,* and *Tgfb1*) in eWAT. **(F)** Relative mRNA expression levels of pro-inflammatory genes (*Tnf* and *Ccl2*) in eWAT. **(G)** Representative H&E and F4/80 immunostaining of eWAT sections from ND- and HFD-fed mice with or without pirfenidone treatment. Scale bars, 100 μm. **(H)** Quantification of adipocyte size in eWAT. **(I)** Quantification of the F4/80-positive area in eWAT. Data are presented as the mean ± SEM (n = 5 mice per group). Statistical significance was determined by two-way ANOVA followed by Tukey’s multiple-comparison test. *P < 0.05; **P < 0.01; ***P < 0.001; ****P < 0.0001; ns, not significant.

### Pirfenidone inhibits fibrogenic ECM programs through the SMAD3–Hippo/YAP axis in vitro and in vivo

To determine how pirfenidone reduces visceral adipose fibrosis, we examined the underlying signaling pathways. First, we established in vitro pro-fibrotic conditions by exposing mature 3T3-L1 adipocytes to transforming growth factor-β1 (TGF-β1). Under these conditions, pirfenidone suppressed induction of extracellular matrix-related and pro-fibrotic transcripts, including *Col1a1*, *Col3a1*, *Fn1*, *Hif1α*, *Lox*, and *Tgfb1* (Fig. 5A). Consistently, pirfenidone reduced the abundance of fibrosis-associated proteins, including α-SMA, FN, and collagen VI (Fig. 5B,C). Given the central role of TGF-β signaling in adipose fibrogenesis and the reported involvement of YAP in matrix-driven transcriptional programs, we examined whether pirfenidone modulates SMAD3 and YAP signaling under pro-fibrotic conditions in adipocytes. Mechanistically, pirfenidone attenuated TGF-β signaling output, as evidenced by decreased SMAD3 phosphorylation and YAP abundance, accompanied by diminished nuclear accumulation of YAP (Fig. 5D,E). These signaling changes were recapitulated in eWAT from HFD-fed mice treated with pirfenidone, which displayed reduced phosphorylated SMAD3 (p-SMAD3) and lower nuclear YAP levels (Fig. 5F,G). Notably, pirfenidone also diminished the phosphorylation of S6K1 and S6, indicative of reduced mTORC1 signaling in adipose tissue (Fig. 5F,H). These coordinated reductions in p-SMAD3, YAP, and mTORC1 output suggest that pirfenidone prevents adipose fibrogenesis in vivo through convergence on the SMAD3–YAP–mTORC1 signaling axis. Together, these results support that pirfenidone suppresses adipose fibrotic dysfunction by dampening TGF-β/SMAD3 signaling and the Hippo effector YAP, thereby limiting adipose-associated ECM gene programs in vitro and in vivo.

**Figure 5.**
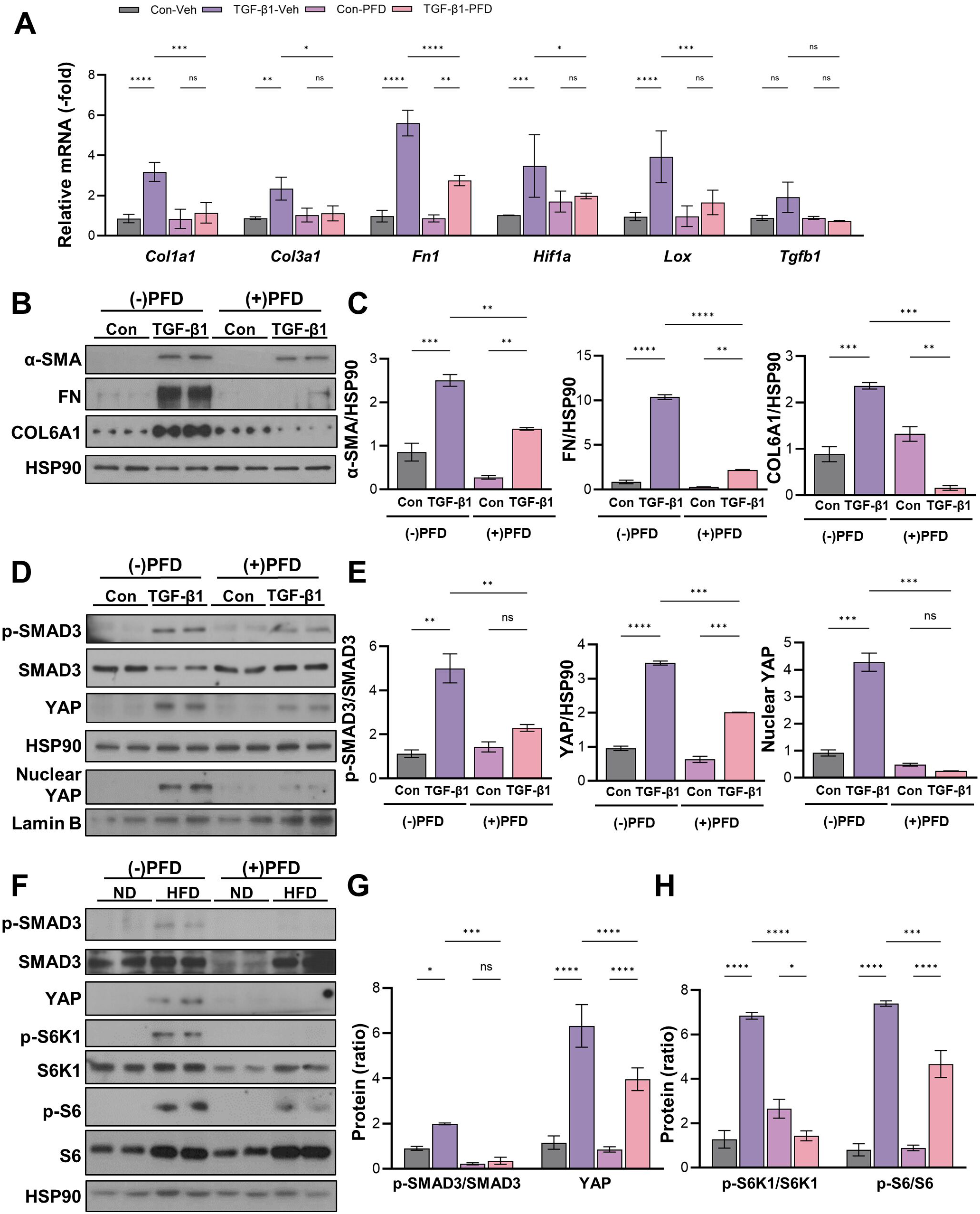
Pirfenidone inhibits ECM deposition via regulating the Hippo pathway. **(A)** Relative mRNA expression levels of extracellular matrix (ECM) and fibrosis-related genes (*Col1a1, Col3a1, Fn1, Hif1α, Lox*, and *Tgfb1*) in differentiated 3T3-L1 adipocytes treated with transforming growth factor-β1 (TGF-β1) in the presence or absence of pirfenidone (PFD). **(B)** Immunoblot analysis of fibrosis-associated proteins α-smooth muscle actin (α-SMA), fibronectin (FN), and collagen VI α1 (Col6A1) in adipocytes treated with TGF-β1 with or without PFD, with quantification normalized to heat shock protein 90 (HSP90). **(C)** Quantification of the α-SMA/HSP90, FN/HSP90, and COL6A1/HSP90 ratios in adipocytes treated with TGF-β1 with or without PFD. **(D)** Immunoblot analysis of phosphorylated mothers against decapentaplegic homolog 3 (p-SMAD3), SMAD3, and yes-associated protein (YAP) in whole-cell lysates (HSP90 as the loading control) and nuclear YAP levels in nuclear fractions (Lamin B as the nuclear loading control). **(E)** Quantification of the p-SMAD3/SMAD3 ratio, the YAP/HSP90 ratio, and nuclear YAP abundance in adipocytes treated with TGF-β1 with or without PFD. **(F)** Immunoblot analysis of phosphorylated SMAD3 (p-SMAD3), total SMAD3, YAP, phosphorylated S6 kinase 1 (p-S6K1), total S6K1, phosphorylated S6 (p-S6), and total S6 in epididymal white adipose tissue (eWAT) from normal diet (ND)- and high-fat diet (HFD)-fed mice treated with vehicle or PFD. Glyceraldehyde-3-phosphate dehydrogenase (GAPDH) was used as a loading control. **(GH)** Densitometric quantification of p-SMAD3/SMAD3, YAP, p-S6K1/S6K1, and p-S6/S6 protein levels normalized to GAPDH and expressed relative to the ND-vehicle group. Data are presented as mean ± SEM (n = 5 mice per group). Statistical significance was determined by two-way ANOVA followed by Tukey’s multiple-comparison test. *P < 0.05; **P < 0.01; ***P < 0.001; ****P < 0.0001; ns, not significant.

### Pirfenidone ameliorates hepatic steatosis and fibrotic injury, coinciding with reduced hepatic mTORC1 output

We next assessed whether pirfenidone improves hepatic pathology in the context of HFD-induced MASLD. Histological analyses showed that pirfenidone reduced hepatic lipid accumulation, as demonstrated by improved H&E morphology and diminished Oil Red O staining (Fig. 6A,B). Consistently, hepatic triglyceride content was significantly decreased in pirfenidone-treated mice (P < 0.05) (Fig. 6C). Beyond steatosis, pirfenidone attenuated fibrotic remodeling, with reduced collagen deposition on Masson’s trichrome and Sirius Red staining (Fig. 6A,D,E). Consistent with these histological and signaling changes, pirfenidone downregulated hepatic lipogenic gene expression (*Fasn*, *Scd1*), inflammatory mediators (*Ccl2*, *Il1b*), and fibrotic gene programs (*Col1a1*, *Col3a1*), while modulating lipid oxidation (Fig. 6F). At the protein level, pirfenidone reduced expression of fibrosis-associated markers (Fig. 6G,H). Notably, pirfenidone suppressed hepatic mTORC1 signaling, as indicated by decreased phosphorylation of both S6K1 and S6 (Fig. 6G,I). Consistent with this in vivo target engagement, pirfenidone reduced S6 phosphorylation in AML12 hepatocytes in a time-and dose-response experiment (EV1 A,B). In addition, pirfenidone attenuated oleic acid–induced neutral lipid accumulation, as measured by boron-dipyrromethene (BODIPY) staining (EV 1C), in parallel with reduced p-S6 levels under lipid-loading conditions (EV 1D). Together, these findings show that pirfenidone improves hepatic steatosis and dampens inflammatory and fibrotic responses in obese MASLD mice, accompanied by reduced activation of mTORC1 output.

**Figure 6.**
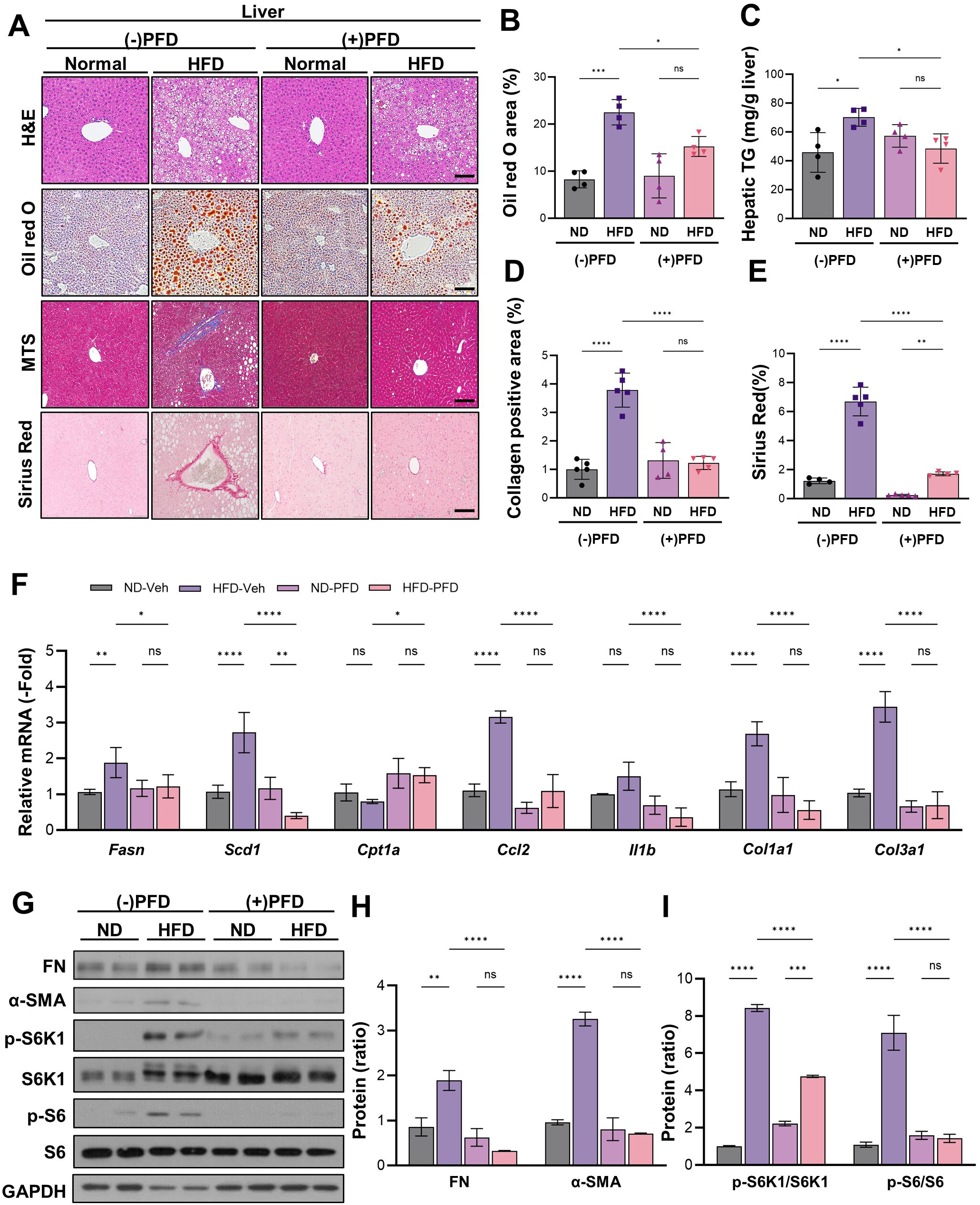
Pirfenidone suppresses hepatic steatosis and fibrosis through inhibition of mTORC1 signaling in the livers of HFD-fed mice. **(A)** Representative histological analyses of liver sections from normal diet (ND)- and high-fat diet (HFD)-fed mice treated with vehicle or pirfenidone (PFD), including hematoxylin and eosin (H&E), Oil Red O, Masson’s trichrome (MTS), and Sirius Red staining. Scale bars, 100 μm. **(B)** Quantification of the Oil Red O–positive area in liver sections. **(C)** Hepatic triglyceride (TG) content in liver tissue. **(D–E)** Quantification of collagen-positive areas based on MTS (D) and Sirius Red (E) staining. **(F)** Relative mRNA expression levels of *Fasn*, *Scd1*, *Cpt1a*, *Ccl2, Il1b*, *Col1a1,* and *Col3a1* in liver tissue. **(G)** Immunoblot analysis of fibronectin (FN), α-smooth muscle actin (α-SMA), phosphorylated S6 kinase 1 (p-S6K1), total S6K1, phosphorylated S6 (p-S6), and total S6 in liver lysates, with glyceraldehyde-3-phosphate dehydrogenase (GAPDH) as a loading control. **(H)** Quantification of FN and α-SMA protein levels normalized to GAPDH. **(I)** Densitometric quantification of p-S6K1/S6K1 and p-S6/S6 ratios in liver tissue. Data are presented as the mean ± SEM (n = 5 mice per group). Each dot represents an individual mouse. Statistical significance was determined by two-way ANOVA followed by Tukey’s multiple-comparison test. *P < 0.05; **P < 0.01; ***P < 0.001; ****P < 0.0001; ns, not significant.

### Pirfenidone blunts adipocyte-derived paracrine signaling to attenuate hepatocyte lipogenic, inflammatory, and fibrogenic programs

To functionally link adipocyte fibrotic remodeling to hepatocyte responses, we established a conditioned medium–based indirect co-culture system in which mature adipocytes were stimulated with TGF-β1 in the presence of vehicle or pirfenidone (Fig. 7A). Conditioned medium from TGF-β1–treated adipocytes induced the expression of lipogenesis-related genes, which was reduced when adipocytes were exposed to pirfenidone (Fig. 7B). In parallel, TGF-β1 adipocyte-conditioned medium increased expression of inflammatory (*Ccl2* and *Tnf*) and fibrosis-associated genes (*Col1a1* and *Col3a1*), whereas pirfenidone-conditioned medium blunted these responses (Fig. 7C). Consistent with a shift toward oxidative metabolism, pirfenidone-conditioned medium increased hepatic expression of fatty acid oxidation–associated genes compared with conditioned medium from fibrotic adipocytes (Fig. 7D). Mechanistically, fibrotic adipocyte-conditioned medium upregulated phosphorylation of inhibitor κB⍺ (IκB⍺) in AML12 cells, consistent with activation of an nuclear factor-κB (NF-κB)-linked inflammatory signaling output, and promoted a fibrogenic protein signature, both of which were attenuated by pirfenidone-conditioned medium (Fig. 7E,F). Collectively, these data support an adipocyte-to-hepatocyte paracrine axis in which pirfenidone dampens hepatocyte lipogenic, inflammatory, and fibrogenic programs triggered by fibrotic adipocytes.

**Figure 7.**
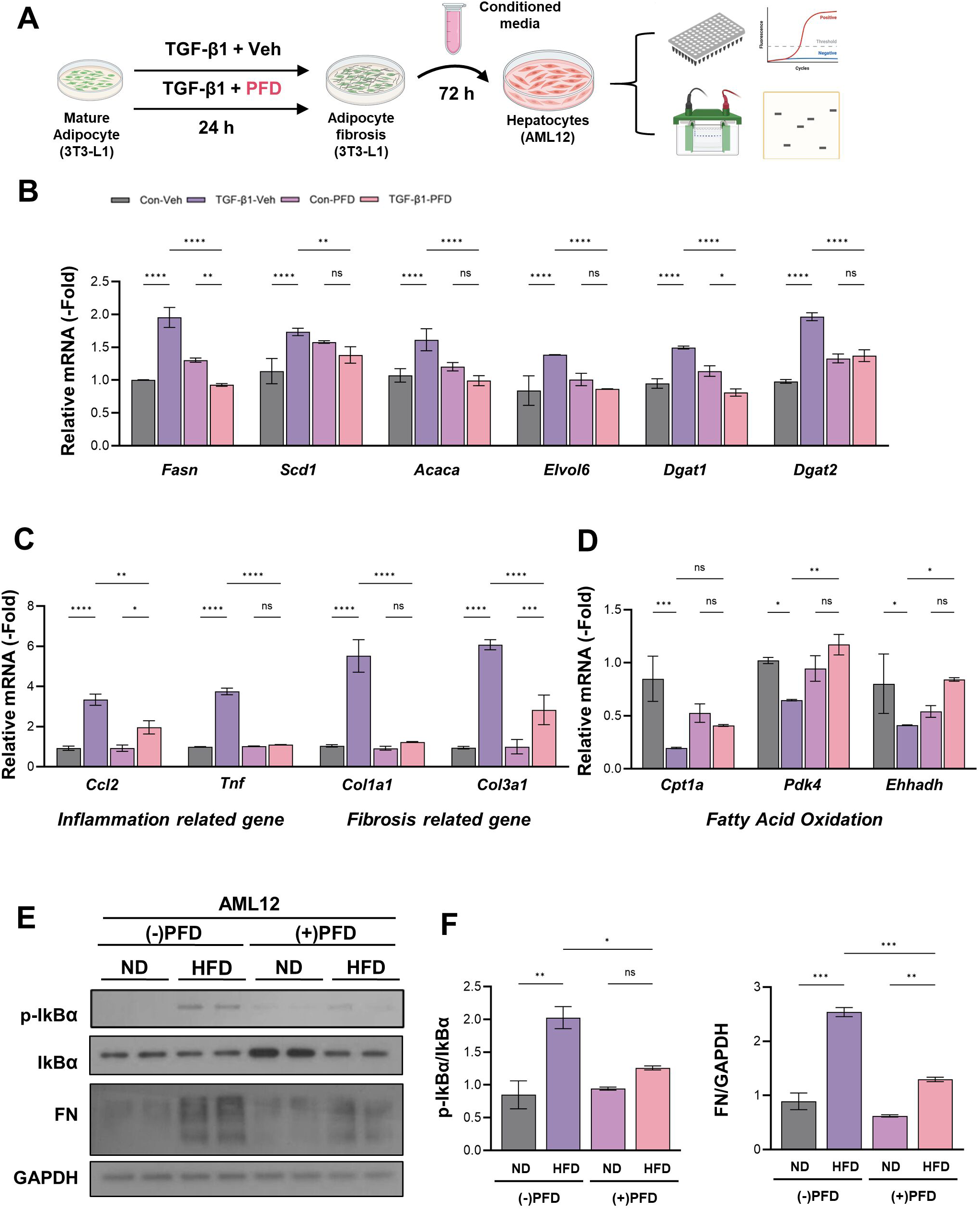
Pirfenidone alleviates adipocyte fibrosis to protect hepatocytes from inflammatory and pro-fibrotic responses through adipocyte–hepatocyte crosstalk. **(A)** Schematic illustration of the adipocyte–hepatocyte co-culture system. Differentiated 3T3-L1 adipocytes were treated with transforming growth factor-β1 (TGF-β1) in the presence or absence of pirfenidone (PFD) to induce a fibrotic adipocyte state. Conditioned medium was collected after 24 h and applied to AML12 hepatocytes for 72 h prior to downstream analyses. **(B)** Relative mRNA expression levels of *Fasn, Scd1, Acaca, Elvol6, Dgat1*, and *Dgat2* in AML12 hepatocytes treated with adipocyte-conditioned medium. **(C)** Relative mRNA expression levels of *Ccl2*, *Tnf*, *Col1a1*, *Col3a1*) in AML12 hepatocytes treated with adipocyte-conditioned medium. **(D)** Relative mRNA expression levels of *Cpt1a, Pdk4*, and *Ehhadh* in AML12 hepatocytes treated with adipocyte-conditioned medium **(E)** Immunoblot analysis of phosphorylated IκBα (p-IκBα), total IκBα, and fibronectin (FN) in AML12 hepatocytes exposed to adipocyte-conditioned medium, with glyceraldehyde-3-phosphate dehydrogenase (GAPDH) as a loading control. **(F)** Quantification of p-IκBα/IκBα ratio and fibronectin protein levels normalized to GAPDH in AML12 hepatocytes. Data are presented as the mean ± SEM. Each dot represents an independent biological replicate. Statistical significance was determined by one-way ANOVA with Tukey’s multiple-comparison test. *P < 0.05, **P < 0.01, ***P < 0.001; ns, not significant.

## Discussion

MASLD/MASH is increasingly recognized as a systemic disorder shaped by maladaptive communication among metabolic organs rather than a liver-centric process (Schuster *et al*, 2018; Targher *et al*, 2021). In particular, depot-specific adipose remodeling can amplify lipid overflow and inflammatory cues that influence hepatocyte metabolism and injury (Azzu *et al*., 2020; Fabbrini *et al*, 2008; Lee *et al*, 2023). Building on prior evidence that pirfenidone ameliorates obesity-associated organ pathology and metabolic dysfunction in experimental steatohepatitis, we asked whether pirfenidone might also target adipose tissue depots, an upstream determinant of metabolic stress, and thereby attenuate adipose–liver crosstalk(Gutierrez-Cuevas *et al*, 2021; Sandoval-Rodriguez *et al*, 2020). In this study, we demonstrate depot-specific adipose effects of pirfenidone in a therapeutic intervention setting, after obesity and MASLD/MASH-like features were established. We show that pirfenidone improves MASLD/MASH-relevant phenotypes by inducing depot-selective adipose tissue remodeling, characterized by attenuated fibro-inflammatory remodeling in eWAT and reduced lipid loading in iWAT.

Using a regimen of 8 weeks of HFD followed by 4 weeks of HFD plus pirfenidone (150 mg/kg), we observed improvements in systemic metabolic parameters, including lower circulating NEFA and ALT levels and improved glucose tolerance and insulin responsiveness (Fig. 1). Importantly, the effects of pirfenidone were not limited to changes in overall adiposity but extended to adipose tissue quality, including adipocyte hypertrophy, inflammatory response, and ECM remodeling, which are associated with metabolic risk and inter-organ stress signaling in obesity.

In subcutaneous iWAT, pirfenidone reduced adipocyte hypertrophy and lipid storage features, consistent with reduced lipid loading (Fig. 2A–C). This was accompanied by suppression of lipogenic gene expression and induction of fatty acid oxidation–associated genes (Fig. 2D,E). Notably, pirfenidone also increased a browning/thermogenic signature that included *Ucp1* and *Ppargc1a* at the mRNA level and PGC1α at the protein level, supporting enhanced mitochondrial oxidative capacity (Fig. 2F,G). Taken together, these changes suggest that pirfenidone shifts iWAT away from lipid storage and toward oxidative lipid utilization and thermogenic programming under obese conditions. This remodeling is consistent with reduced lipid overflow and improved systemic metabolic profiles.

In visceral eWAT, pirfenidone improved fibro-inflammatory remodeling despite minimal change in depot mass. Collagen deposition was reduced, as assessed by Masson’s trichrome and Sirius Red staining. This histological improvement was corroborated by decreased levels of fibrosis-associated proteins (α-SMA, FN, and collagen VI) and broad suppression of a pro-fibrotic transcriptional program (Fig. 4A–E). Pirfenidone also decreased inflammatory mediators, such as *Tnf* and *Ccl2*, and reduced macrophage accumulation, which was accompanied by decreased adipocyte hypertrophy (Fig. 4F–I). These findings suggest that pirfenidone suppresses fibrotic ECM remodeling in visceral adipose tissue, with concurrent attenuation of inflammatory activation in obesity-associated metabolic dysfunction.

This interpretation is supported by clinical and experimental evidence linking adipose fibrogenesis to MASLD. In humans, increased adipose tissue fibrogenesis is associated with non-alcoholic fatty liver disease in obesity even when adipose expandability is not impaired, suggesting that adipose ECM remodeling is a clinically relevant feature of metabolically unhealthy obesity (Beals *et al*., 2021). Mechanistic studies in mice suggest that adipose matrix remodeling can drive systemic metabolic dysfunction and hepatic steatosis through collagen-derived mediators (Sun *et al*., 2014). These observations align with an adipocentric view of MASLD in which adipose tissue failure promotes lipid overflow and pathological inter-organ crosstalk (Lee *et al*., 2023).

Mechanistically, our data indicate that pirfenidone suppresses fibrogenic signaling outputs through coordinated modulation of TGF-β/SMAD3 and the Hippo effector YAP in eWAT. In a pro-fibrotic adipocyte setting induced by TGF-β1, pirfenidone reduced ECM-related transcripts and fibrosis-associated proteins while attenuating SMAD3 phosphorylation (Fig. 5A–E). Pirfenidone also decreased YAP levels and nuclear accumulation, consistent with reduced transcriptional outputs that reinforce matrix programs. Importantly, reduced p-SMAD3 and YAP levels were recapitulated in eWAT from HFD-fed mice treated with pirfenidone, supporting in vivo relevance (Fig. 5F,G). Collectively, reduced SMAD3 phosphorylation and YAP nuclear accumulation were associated with broad suppression of ECM programs. However, further studies will be required to clarify the mechanistic relationships among these nodes and to determine whether pirfenidone primarily disrupts upstream pro-fibrotic cues in eWAT.

Across metabolic tissues, pirfenidone was associated with reduced mTORC1 output, as indicated by decreased phosphorylation of S6K1 and S6 in vitro and in vivo. Stage-specific exposure to pirfenidone during adipocyte differentiation attenuated mTORC1 signaling and delayed neutral lipid accumulation, with corresponding reductions in lipogenic and lipid-uptake gene expression (Fig. 3E,G,H). These findings indicate a direct, adipocyte-autonomous effect of pirfenidone in constraining mTORC1-dependent anabolic programs during adipogenesis (Song *et al*, 2023; Zhou *et al*, 2023). In contrast, fatty acid oxidation–related transcripts were not significantly altered in vitro, suggesting that pirfenidone does not directly promote oxidative metabolism in adipocytes (Fig. 3D). The increased fatty acid oxidation observed in vivo is therefore more likely to reflect secondary adaptations within the tissue microenvironment, potentially involving systemic or intercellular cues (Liu *et al*, 2022; Tsukada *et al*, 2023). However, the precise mechanisms underlying these context-dependent effects remain to be elucidated. Pirfenidone treatment also reduced AMPK phosphorylation, suggesting that it alters lipid-associated programs by inhibiting mTORC1 signaling rather than activating AMPK (Fig. 3E) (Lee *et al*, 2020; Lee *et al*, 2018). In eWAT, the inhibited mTORC1 pathway occurred in parallel with diminished SMAD3–YAP signaling and ECM programs, suggesting that mTORC1 may amplify fibrogenic remodeling in this depot (Fig. 5F–H). Given the broad role of mTORC1 in anabolic metabolism and stress-responsive programs (Adegoke *et al*, 2012), we interpret mTORC1 suppression as a component of pirfenidone action consistent with reduced lipogenic/storage programs and improved adipose tissue architecture rather than as a single linear driver of all downstream effects.

A major advance of this study is the functional demonstration of an adipocyte-to-hepatocyte signaling axis linking adipocyte fibrotic remodeling to hepatocyte stress responses. Conditioned medium from TGF-β1–treated mature adipocytes induced hepatocyte lipogenic gene expression and increased inflammatory and fibrogenic programs in AML12 hepatocytes (Fig. 7). When adipocytes were treated with pirfenidone during pro-fibrotic stimulation, the conditioned medium elicited attenuation of hepatocyte lipogenic, inflammatory, and fibrogenic responses and was associated with increased expression of fatty acid oxidation–related genes. These findings demonstrate that suppressing a fibrogenic adipocyte state can translate into reduced hepatocyte metabolic stress programs, suggesting that adipose tissue quality is a key upstream determinant of liver injury pathways.

In parallel with adipose remodeling and crosstalk effects, pirfenidone improved hepatic steatosis and reduced inflammatory and fibrotic markers in HFD-fed mice (Fig. 6). Multiple readouts suggested reduction of lipid accumulation and attenuation of fibrotic remodeling. These improvements coincided with reduced hepatic mTORC1 output (p-S6K1 and p-S6). In hepatocytes, pirfenidone reduced p-S6 in time- and dose-response experiments and attenuated oleic acid–induced lipid accumulation in AML12 cells (EV 1), supporting hepatocyte target engagement, consistent with the in vivo findings. Together, our data suggest that pirfenidone may act through both adipose-mediated and direct hepatic mechanisms, thereby improving hepatic steatosis and injury.

This study has limitations that can be addressed in future work. First, although our conditioned-medium system provides functional evidence for adipocyte-to-hepatocyte communication, direct co-culture approaches will help define the contributions of adipose-mediated versus hepatocyte-autonomous mechanisms. Second, although our data implicate the SMAD3–YAP signaling axis in adipose fibrogenic remodeling, genetic or pharmacologic approaches (e.g., modulation of YAP/TAZ activity, SMAD3 inhibition, or mTORC1 rescue) will be required to establish causality and determine the functional order among these nodes. Third, the cause of depot specificity remains unknow and may involve differences in stromal composition, mechanical constraints, immune cell dynamics, and the nature of local pro-fibrotic cues.

In summary, our findings elucidate the adipose actions of pirfenidone and establish depot-selective adipose remodeling as a mechanistic bridge between obesity-associated adipose dysfunction and hepatic pathology. By restraining lipid accumulation in iWAT and suppressing fibro-inflammatory remodeling in eWAT through dampened mTORC1 output and reduced SMAD3–YAP signaling, pirfenidone alters adipose tissue quality and adipose–liver communication. These results underscore the importance of targeting maladaptive inter-organ crosstalk and suggest pirfenidone as a potential therapeutic for obesity-associated MASLD/MASH.

## Methods

### Animal experiments

Male C57BL/6J mice were obtained from Japan SLC, Inc. Mice were maintained at 22 ± 2°C with ad libitum access to food and water under a 12 h light/dark cycle. To evaluate pirfenidone in vivo, mice were fed an HFD for 12 weeks. Pirfenidone (150 mg/kg) or vehicle was administered by oral gavage three times per week starting at week 9 and continued until the experimental endpoint. At sacrifice, serum, eWAT, iWAT, and liver tissues were collected for analysis.

### Oral glucose tolerance test and insulin tolerance test

For the oral glucose tolerance test (OGTT), mice were fasted for 4 h and administered glucose (1 g/kg body weight) by oral gavage. For the insulin tolerance test (ITT), mice were fasted for 4 h and injected intraperitoneally with insulin (0.75 U/kg body weight), followed by serial glucose measurements at indicated time points.

### Serum biochemistry

Serum alanine transaminase (ALT) levels were measured using FUJI DRI-CHEM SLIDE kits and analyzed on a colorimetric analyzer according to the manufacturer’s instructions. Serum non-esterified fatty acid (NEFA) levels were measured using a commercial colorimetric assay kit (Wako) according to the manufacturer’s instructions.

### Histology and immunostaining

Tissues were fixed in 10% neutral-buffered formalin overnight at 4°C. For paraffin sections, tissues were deparaffinized, rehydrated, and processed for antigen retrieval prior to immunohistochemistry. H&E, Masson’s trichrome, and Sirius Red staining were performed using standard protocols. Images were acquired using a Nikon Coolscope microscope and analyzed using ImageJ. Quantification was performed in a blinded manner using identical threshold settings across groups.

### Hepatic steatosis measurement

Oil Red O staining was performed on liver cryosections, followed by Mayer’s hematoxylin counterstaining. Briefly, liver tissues were embedded in O.C.T compound and sectioned at 5-µm thickness. Cryosections were air-dried at room temperature for at least 30 min and fixed in 10% formalin for 5 min. Sections were then dehydrated in 100% propylene glycol for 5 min and stained with Oil Red O solution at 60°C for 10 min. Excess stain was removed with 85% propylene glycol, and sections were counterstained with Mayer’s hematoxylin for 1 min to visualize nuclei. Slides were mounted using an aqueous mounting medium and analyzed under a microscope within 2 h of preparation. Lipid-positive areas were quantified in a blinded manner using identical threshold settings from three non-overlapping fields per section.

### Cell culture and adipocyte differentiation

AML12 hepatocytes were cultured in Dulbecco’s Modified Eagle’s Medium (DMEM) supplemented with 10% fetal bovine serum (FBS) and 1% penicillin/streptomycin at 37°C in a humidified atmosphere with 5% CO₂. The 3T3-L1 preadipocytes were kindly provided by Dr. J. Kim (Yonsei University College of Medicine) and maintained in DMEM supplemented with 10% calf serum and 1% penicillin/streptomycin at 37°C in a humidified 5% CO₂ atmosphere. For adipocyte differentiation, post-confluent cells were induced (day 0) with differentiation medium (DMEM + 10% FBS) containing an MDI cocktail—3-isobutyl-1-methylxanthine (IBMX, 520 µM), dexamethasone (1 µM), and insulin (1 µg/mL)—for 48 h. Cells were then maintained in DMEM + 10% FBS with insulin for 48 h, followed by DMEM + 10% FBS with medium changes every 48 h until day 8. Fully differentiated adipocytes were used for subsequent assays. To induce a pro-fibrotic response, differentiated 3T3-L1 adipocytes were treated with TGF-β1 (10 ng/mL) in the presence of pirfenidone (500 µM) or vehicle (dimethyl sulfoxide, DMSO) for 24 h before harvest.

### Pro-fibrotic adipocyte setting

To assess adipocyte-to-hepatocyte signaling, conditioned medium was generated by treating fully differentiated 3T3-L1 adipocytes with TGF-β1 (10 ng/mL) ± pirfenidone (500 µM). Conditioned medium was applied to AML12 hepatocytes for 72 h, after which cells were harvested for downstream analyses.

### Oil Red O staining

For Oil Red O staining of cultured cells, a working solution was prepared by mixing 0.5% Oil Red O in isopropanol with deionized water at a 3:2 ratio. On day 8 of differentiation, cells were washed with Dulbecco’s phosphate-buffered saline (DPBS), fixed with 4% formaldehyde for 15 min at room temperature, and stained with the Oil Red O working solution for 1 h. Cells were washed thoroughly with deionized water to remove excess dye. For quantitative analysis, retained dye was eluted with isopropanol, and absorbance was measured at 500 nm using a microplate reader (VERSA Max).

### Immunoblotting

Cells and tissues were lysed in buffer containing 50 mM Tris–HCl (pH 7.5), 150 mM NaCl, 1% Nonidet P-40 (NP-40), and protease inhibitors. Lysates were clarified by centrifugation (20,000 × g, 15 min, 4°C), and protein concentrations were determined by Bradford assay. Equal amounts of protein were resolved by sodium dodecyl sulfate polyacrylamide gel electrophoresis (SDS–PAGE), transferred to polyvinylidene difluoride (PVDF) membranes, and blocked in 5% skim milk for 1 h. Membranes were incubated with primary antibodies overnight at 4°C and horseradish peroxidase (HRP)-conjugated secondary antibodies for 1 h at room temperature. Signals were detected using enhanced chemiluminescence (ECL) and quantified within the linear range.

### RNA isolation and quantitative RT–PCR

Total RNA was extracted using TRIzol and reverse-transcribed into cDNA using a commercial cDNA synthesis kit. Quantitative PCR was performed using SYBR Green chemistry on an ABI PRISM 7700 system. Gene expression was normalized to 18S rRNA. The primer sequences are provided in Supplementary Table 1.

### Data analysis, statistics, and reproducibility

All experiments were performed with at least two independent biological replicates unless stated otherwise. Independent experiments were conducted by different investigators to ensure reproducibility. Animals were allocated to experimental groups based on genotype and dietary intervention. No samples or animals were excluded from the analyses unless required for animal welfare reasons. Quantification and data analysis were performed using predefined criteria applied uniformly across experimental groups. The sample size (n) refers to biologically independent samples or animals, as indicated in the figure legends.

Data are presented as the mean ± standard error of the mean (SEM) unless otherwise stated. When the number of biological replicates was less than five, individual data points are shown alongside summary statistics. Statistical analyses were performed using GraphPad Prism (v10.2.2). Comparisons between two groups were conducted using two-tailed unpaired Student’s t-tests, and comparisons among multiple groups were performed using one-way or two-way analysis of variance (ANOVA) followed by Tukey’s post hoc multiple-comparison test, as specified in the figure legends. A P value < 0.05 was considered statistically significant.

### Study approval

All animal experiments were approved by the Animal Care and Use Committee of Yonsei University College of Medicine (approval numbers: 2020-0274, 2022-0345, and 2023-0161) and conducted in accordance with institutional guidelines and ARRIVE 2.0.

## Data Availability

All data supporting the findings of this study are included in the article and its Expanded View figures, and are available from the corresponding author upon reasonable request.

## Acknowledgements

We thank the members of our research team at Yonsei University College of Medicine and the Korea University College of Pharmacy for their technical assistance. The graphical abstract was created using BioRender. The authors and their funder had full access to all data in this study and take full responsibility for the decision to submit this article.

## Author contributions

Yu Seol Lee designed the research and acquired the data. Yu Seol Lee, Ji Yun Bang, Shin Young Cha, and Eo Jin Lee analyzed the data. Yu Seol Lee and Ji Yun Bang wrote the manuscript. Yu Seol Lee, Ji Yun Bang, Da Hyun Lee, Da Ye Kim, and Jisu Han performed the in vivo experiments. Soo Han Bae and Yu Seol Lee conceptualized the study, contributed to the manuscript revision, and supervised the project. All authors critically reviewed the manuscript drafts and approved the final manuscript.

## Ethics declarations

All procedures performed in studies involving human participants were in accordance with the ethical standards of the institutional and/or national research committee and with the 1964 Helsinki Declaration and its later amendments or comparable ethical standards.

## Disclosure and competing interests statement

The authors declare that they have no conflicts of interest.

### The Paper Explained

#### PROBLEM

Obesity-driven MASLD/MASH progression is closely linked to adipose tissue dysfunction, acute expansion, and fibrogenesis. These abnormalities are increasingly recognized as key drivers of systemic metabolic disorder through adipose–liver crosstalk. Although pirfenidone is clinically approved as an anti-fibrotic agent, its contribution to modulating adipose–liver communication in metabolic liver disease remains poorly defined.

#### RESULTS

Our study reveals that pirfenidone plays a critical role in MASH progression by modulating the adipose tissue–liver axis. Pharmacological injection of pirfenidone alleviates hepatic steatosis, inflammation, and fibrosis in mouse models while reducing adipose tissue dysfunction. These effects are achieved through inhibition of the mTORC1 pathway. Mechanistically, pirfenidone blunts pro-lipogenic and pro-fibrotic signaling in AML12 cells induced by conditioned medium from fibrotic adipocytes.

#### IMPACT

Pirfenidone emerges as a promising therapeutic for MASH, addressing both hepatic and adipose tissue dysfunctions. Targeting adipose tissue remodeling within the adipose tissue–liver axis may lead to effective treatments in patients with obesity-associated MASH.

## Abbreviations

eWAT: epididymal white adipose tissue
HFD: high-fat diet
iWAT: inguinal white adipose tissue
MASH: metabolic dysfunction–associated steatohepatitis
MASLD: metabolic dysfunction–associated steatotic liver disease
mTORC1: mechanistic target of rapamycin complex 1
SAT: subcutaneous adipose tissue
VAT: visceral adipose tissue

